# SHAMAN: a user-friendly website for metataxonomic analysis from raw reads to statistical analysis

**DOI:** 10.1101/2019.12.18.880773

**Authors:** Stevenn Volant, Pierre Lechat, Perrine Woringer, Laurence Motreff, Christophe Malabat, Sean Kennedy, Amine Ghozlane

**Affiliations:** Hub de Bioinformatique et Biostatistique – Département Biologie Computationnelle, Institut Pasteur, USR 3756 CNRS, Paris, France; Biomics – Département Génomes et Génétique, Institut Pasteur, Paris, France

**Keywords:** Metagenomics, Statistical analysis, Visualization

## Abstract

Comparing the composition of microbial communities among groups of interest (e.g., patients vs healthy individuals) is a central aspect in microbiome research. It typically involves sequencing, data processing, statistical analysis and graphical representation of the detected signatures. Such an analysis is normally obtained by using a set of different applications that require specific expertise for installation, data processing and in some case, programming skills. Here, we present SHAMAN, an interactive web application we developed in order to facilitate the use of (i) a bioinformatic workflow for metataxonomic analysis, (ii) a reliable statistical modelling and (iii) to provide among the largest panels of interactive visualizations as compared to the other options that are currently available. SHAMAN is specifically designed for non-expert users who may benefit from using an integrated version of the different analytic steps underlying a proper metagenomic analysis. The application is freely accessible at http://shaman.pasteur.fr/, and may also work as a standalone application with a Docker container (aghozlane/shaman), conda and R. The source code is written in R and is available at https://github.com/aghozlane/shaman. Using two datasets (a mock community sequencing and published 16S metagenomic data), we illustrate the strengths of SHAMAN in quickly performing a complete metataxonomic analysis.

## Introduction

Quantitative metagenomic techniques have been broadly deployed to identify associations between microbiome and environmental or individual factors (e.g., disease, geographical origin, etc.). Analyzing changes in the composition and/or the abundance of microbial communities yielded promising biomarkers, notably associated with liver cirrhosis(1), diarrhea(2), colorectal cancer(3), or associated with various pathogenic(4) or probiotic effects(5) on the host.

In metataxonomic studies, a choice is made prior to sequencing in order to specifically amplify one or several regions of the rRNA (usually the 16S or the 18S rRNA for procary-otes/archaea and the ITS, the 23S or the 28S rRNA for eukaryotes) so that the composition of microbial communities may be characterized with affordable techniques.

A typical workflow includes successive steps: (i) OTU (Operational Taxonomic Unit) picking (dereplication, denoising, chimera filtering and clustering)(6), (ii) OTU quantification in each sample and (iii) OTU annotating with respect to a reference taxonomic database. This process may require substantial computational resources depending on both the number of samples involved and the sequencing depth. Several methods are currently available to complete these tasks, such as Mothur(7), Usearch(8), DADA2(9) or Vsearch(10). The popular Qiime(11) simplifies these tasks (i to iii) and visualizations by providing a python-integrated environment. Schematically, once data processing is over, both a contingency table and a taxonomic table are obtained. They contain the abundance of OTUs in the different samples and the taxonomic annotations of OTUs, respectively. The data are normally represented in the standard BIOM format(12).

Statistical analysis is then performed to screen significant variation in microbial abundance. To this purpose, several R packages were developed, such as Metastats(13) or Metagenomeseq(14). It is worth noticing that other approaches which were originally designed for RNA-seq, namely DESeq2(15) and EdgeR(16), are also commonly used to carry out metataxonomic studies(17, 18). They provide an R integrated environment for statistical modelling in order to test the effects of a particular factor on OTU abundance. Nevertheless using all of these different methods requires a technical skills in Unix, R and experience in processing metagenomics data. To this end, we developed SHAMAN in order to provide a method that simplifies the analysis of metataxonomic data, especially for users who are not familiar with the technicalities of bioinformatic and statistical methods that are commonly applied in this field.

SHAMAN is an all-inclusive approach to estimate the composition and abundance of OTUs, based on raw sequencing data, and to perform the statistical analysis of the processed files. First, the user can submit raw data in FASTQ format and define the parameters of the bioinformatic workflow. The output returns a BIOM file for each database used as reference for annotation, a phylogenetic tree in Newick format as well as FASTA-formatted sequences of all OTUs that were identified. The second step consists in performing statistical analysis. The user has to provide a “target” file that associates each sample with one or several explanatory variables. These variables are automatically detected in the target file. An automatic filtering of the contingency matrix of OTUs may be activated in order to remove features with low frequency. Setting up the contrasts to be compared was also greatly simplified. It consists in filling in a form that orients the choices of users when defining the groups of interest. Several options to visualize data are available at three important steps of the process: quality control, bio-analysis and contrast comparison. At each step, a number of common visual displays are implemented in SHAMAN to explore data. In addition, SHAMAN also includes a variety of original displays that is not available in other applications such as an abundance tree to visualize count distribution according to the taxonomic tree and variables, or the logit plot to compare feature p-values in two contrasts. Figures may be tuned to emphasize particular statistical results (e.g., displaying significant features in a given contrast, performing intersection between contrasts), to be more specific (e.g. feature abundance in a given modality) or to improve the aesthetics of the graph (by changing visual parameters). Figures fit publication standards and the corresponding file can be easily downloaded.

Several web applications were developed to analyze data of metataxonomic studies, notably, FROGS(19) as well as Qiita(20) for bioinformatic data processing, Shiny-phyloseq(21) for statistical analysis, Metaviz(22) and VAMPS2(23) that make a particular focus on data visualization. While these interfaces propose related functionalities, the main specificity of SHAMAN is to combine of all these steps in a single user-friendly application. Last, SHAMAN may register a complete analysis which may be of particular interest for matters of reproducibility.

## DESCRIPTION

SHAMAN is implemented in R using the shiny-dashboard framework. The application is divided into three main components (Fig. S1): a bioinformatic workflow to process the raw FASTQ-formatted sequences, a statistical workflow to normalize and further analyse data, as well as a visualization platform.

### Metataxonomic pipeline

The bioinformatic workflow implemented in SHAMAN relies on the Galaxy platform(24) that provides modular and scalable analyses. SHAMAN includes a daemon-program (written in Python) using bioblend(25) to communicate with Galaxy. It is worth noticing that previous studies, e.g. performed on mosquito microbiota(26), showed that some non-annotated OTUs turned to be sequences of the host organism. To overcome such issues, the user can optionally filter out reads that align with the host genome and the PhiX174 genome (used as a control in Illumina sequencers). The latter task is performed with Bowtie2 v2.2.6(27). By default, quality of reads is checked with AlienTrimmer(28) v0.4.0, a software for trimming off contaminant sequences and clipping. Paired-end reads are then merged with Pear(29) v0.9.10.1. OTU picking, taxonomic annotation and OTU quantification are performed using Vsearch(10) v2.3.4.0, a software which is both accurate and efficient (6). The process also includes several steps of dereplication, singleton removal and chimera detection. By default, clustering is performed with a threshold of 97% in sequence identity. The input amplicons are aligned against the set of detected OTUs to create a contingency table containing the number of amplicons assigned to each OTU. The taxonomic annotation of OTUs is performed based on various databases, i.e., with SILVA(30) rev. 132 SSU (for 16S, 18S) and LSU (for 23S and 28S sequences), Greengenes(31) (for 16S, 18S sequences) and Underhill rev. 1.6.1(32), Unite rev. 8.0(33) and Findley(34) for ITS sequences. These databases are kept up-to-date every two month with biomaj.pasteur.fr. OTU annotations are filtered according to their identity with the reference(35). Phylum annotations are kept when the identity between the OTU sequence and reference sequence is ≥ 75%, ≥ 78.5% for classes, ≥ 82% for orders, ≥ 86.5% for families, ≥ 94.5% for genera and ≥ 98% for species. In addition, a taxonomic inference made based on a naive Bayesian approach, RDP classifier(36) v2.12, is systematically provided. By default, RDP annotations are included whenever the annotation probability is ≥ 0.5. All the abovementioned thresholds may be tuned by the user.

A phylogenetic analysis of OTUs is provided: multiple alignments are obtained with Mafft(37) v7.273.1, filtering of regions that are insufficiently conserved is processed using BMGE(38) v1.12 and finally, FastTree(39) v2.1.9 is used to infer the phylogenetic tree. Based on the latter tree, a Unifrac distance(40) may be computed in SHAMAN to compare microbial communities. The outcomes of the overall workflow are stored in several files: a BIOM file (per reference database), a phylogenetic tree as well as a summary file describing the number of elements passing the differents steps of the workflow. The data are associated to a key that is unique to a project. Such a key allows to automatically reload all the results previously obtained in a given project.

### Statistical workflow

The statistical analysis in SHAMAN is based on DESeq2 which is a method to model OTU counts with a negative binomial distribution. It is known as one of the most accurate approach to detect differentially abundant bacteria in metagenomic data(17, 18). Relying on a robust estimation of variation in OTUs, the DESeq2 method shows suitable performances with datasets characterized by a relatively low number of observations per group together with a high number of OTUs.

This method typically requires the following input files: a contingency table, a taxonomic table and a target file describing the experimental design. These data are processed to generate a meta-table that assign to each OTU a taxonomic annotation and a raw count per sample.

### Normalization

Normalization of the raw counts is one of the key issues when analyzing microbiome experiments. The uniformity of the sequencing depth is affected by sample preparation and dye effects(41). Normalizing data is therefore expected to increase the accuracy of comparisons. It is done by adjusting the abundance of OTUs across samples. Four different normalization methods are currently implemented in SHAMAN. For the sake of consistency, all of these methods are applied at the OTU level.

A first method is the relative log expression (RLE) normalization and is implemented in the DESeq2 package. It consists in calculating a size factor of each sample, i.e., a multiplication factor that increases or decreases the OTU counts in samples. It is defined as the median ratio, between a given count and the geometric mean of each OTU. Such a normalization was shown to be suited for metataxonomic studies(17). In practice, many OTUs are found in a few samples only, which translate into sparse count matrices(14). In this case, the RLE method may lead to a defective normalization - as only a few OTU are taken into account - or might be impossible if all OTUs show a null abundance in one sample at least. In the Phyloseq(42) R package, the decision was made to replace the null abundance by a count of 1. In SHAMAN, we decided to include two new normalization methods. They are modified versions of the original RLE so that they better account for matrix sparsity (number of zero-valued elements divided by the total number of elements). In the *non-null normalization* (1) cells with null values are excluded from the computation of the geometric mean. This method therefore takes all OTUs into account when estimating the size factor. In the second method that we coined as the *weighted non-null normalization* (2), weights are introduced so that OTUs with a big number of occurrences have a higher influence when calculating the geometric mean.

Assume that *C* = (*c_ij_*)1≤*i*≤*k*;1≤*j*≤*n* is a contingency table where *k* and *n* are the number of features (e.g. OTUs) and the number of samples, respectively. Here, *c_ij_* represents the abundance of the feature *i* in the sample *j*. The size factor of sample *j* is denoted by *s_j_*.

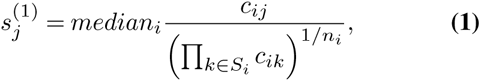

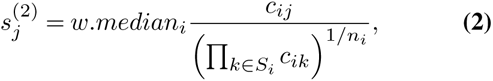

where *S_i_* stands for the subset of samples with non null values for the feature *j* and *n_i_* is the size of with 30 samples; this subset. The function *w.median* corresponds to a weighted median.

An alternative normalization technique is the *total counts*(43) which is convenient for highly unbalanced OTU distribution among samples.

Using a simulation-based approach, we addressed the question of the performance of the *non-null* and the *weighted nonnull normalization* techniques when the matrix sparsity and the number of observations increase. We compared these new methods to those normally performed with DESeq2 and Phyloseq. To do so, we normalized 500 matrices with varied sparsity levels (i.e., 0.28, 0.64 and 0.82) and a different number of observations (i.e., 4, 10 and 30). We calculated the average coefficient of variation (CVmean)(44) for each normalization method (Fig. S2). Considering that these OTUs are assumed to have relatively constant abundance within the simulations, the coefficient of variation is expected to be lower when the normalization is more efficient. Overall, the *non-null* and the *weighted non-null* normalization methods exhibited a lower coefficient of variation as compared to the other methods, when sparsity in the count matrix is high and the number of observations is increased. These differences were clear especially when comparing DESeq2 and Phyloseq to the *weighted non-null normalization* (sparsity ratio of 0.28, 0.64 and 0.82, with 30 samples; t-tests *p* < 0.001) (Fig. S2).

### Contingency table filtering

In metataxonomic studies, contingency tables are often very sparse and after statistical analysis, some significant differences among groups may not be of great relevance. This may arise when a feature, distributed in many samples with a low abundance, is slightly more abundant in a group of comparison. These artifacts are generally excluded by DESeq2 with an independent filtering. Furthermore, if a feature is found in a few samples only, it may lead to non-reliable results when its abundance is high (when it is not 0). Such distributions may also impact the normalization process as well as the dispersion estimates. In order to avoid misinterpretation of results, we propose an optional extra-step of filtering: by excluding features characterized by a low abundance and/or a low number of occurrence in samples (e.g. features occurring in less than 20% of the samples). To set a by-default abundance threshold, SHAMAN search for an inflection point at which the curve between the number of observations and the abundance of feature changes from being linear to concave. This process is performed with linear regression in the following manner:

1. We define *I* the interval 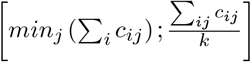.
2. For each *x* ∈ *I*, we compute *h*(*x*) defined as the number of observations with a total abundance higher than *x*.
3. We compute the linear regression between *h*(*x*) and *x*.
4. The intercept is set as the default threshold.

(see Appendix 1 for more information). This extra-filtering aims at refining the first filtering processed with DESeq2 and may lead to a significant decrease of the computation time. The impact of filtering steps may be visually assessed with plots displaying the features that will be included in the analysis and those that will be discarded.

### Statistical modelling

The statistical model relies on the variables that are available in the file of experimental design. By default, all variables are included in the model but the end-user can edit this selection and further add interactions between variables of interest. In addition, other variables such as batches or clinical data (e.g., age, sex, etc.) may be used as covariates. SHAMAN then automatically checks whether the model is statistically suitable (i.e., whether all the model parameters may be estimated). When it is not the case, an warning message appears and a “how to” box proposes a practical way to solve the issue. In SHAMAN, statistical models may be fitted at any taxonomic levels. Normalized counts are summed up within a given a taxonomic level.

To extract features that exhibit significant differential abundance (between two groups), the user must define a contrast vector. Both a guided mode and an expert mode are available in SHAMAN. In the guided mode, the user specifies the groups to be compared using a dropdown menu. This mode is only available for DESeq2 v1.6.3 which is implemented in DESeq2shaman package (https://github.com/aghozlane/DESeq2shaman). In advanced comparisons, the user may define a contrast vector by specifying coefficients (e.g., −1, 0, 1) assigned to each variable.

### Visualization

After running a statistical analysis, many displays are available:

i. Diagnostic plots (such as barplots, boxplots, PCA, PCoA, NMDS and hierarchical clustering) help the user examine both raw and normalized data. For instance, these plots may reveal clusters, sample mislabelling and/or batch effects. Scatterplots of size factors and dispersion (i.e., estimates that are specific to DESeq2) are useful when assessing both the relevance and robustness of statistical models. PCA-and PCoA-plots associated with a PERMANOVA test may be used as preliminary results in the differential analysis as they may reveal global effects among groups of interest.
ii. Significant features are gathered in a table including, the base mean (mean of the normalized counts), the fold change (i.e., the factor by which the average abundance changes from one group to the other), as well as the corresponding adjusted p-values. The user may view tables for any contrasts and can export it into several formats. Volcano plots and bar charts of p-values and log2 fold change are also available this section.
iii. A global visualization section provides a choice of 9 interactive plots such as barplots, heatmaps and boxplots to represent differences in abundance across groups of interest. Diversity plots display the distribution of various diversity indices: alpha, beta, gamma, Shannon, Simpson and inverse Simpson. Scatterplots and network plots show association between feature abundance with other variables from the target file. To explore variations of abundance across the taxonomic classification, we included an interactive abundance tree and a Krona plot(45). Rarefaction curves are of great use to further consider the number of features in samples with respect to the sequencing depth.
iv. In the comparison section, plots displaying comparisons among contrasts may be created. It includes several options such as, Venn diagram or upsetR graph(46) (displaying subsets of common features across contrast), heatmap, a logit plot(47) (showing the log2 fold-change values in each feature), a density plot and a multiple Venn diagram to summarize the number of features captured by each contrast. All these graphs can be exported into four format (eps, png, pdf and svg).

## APPLICATION

### Comparison of SHAMAN with other available tools for meta-taxonomic analyses

A brief qualitative assessment of the strengths and limits of SHAMAN was done in comparison with other similar web interfaces (Table 1). We first identified a list of important considerations that have practical implications for the user such as processing of raw sequencing data, statistical workflow, visualization, data storage and accessibility. For each similar web interface, we then evaluated whether it met those criteria. Besides that SHAMAN presents a number of advantages, we think that such nested solution is essential for a careful interpretation of the results. Any results in SHAMAN may be cross-checked with a quantification or an annotation performed at an earlier stage. Furthermore several applications presented in Table 1, impose the burden to import/export R objects which requires skills in R programming. It may also represent a source of issues for reproducibility, notably in terms of compatibility of the packages over time.

**Table 1.** Comparison of SHAMAN with other web interface for metataxonomic analysis.

| Category | SHAMAN | FROGS | Qiita | Shiny-phyloseq | Metaviz | Vamps |
| --- | --- | --- | --- | --- | --- | --- |
| OTU processing | Yes | Yes | Yes | No | No | No |
| Normalization | Yes | Yes | No | No | No | No |
| Modelisation | Yes | Manova | No | D | M | No |
| Diversity analysis | Yes | Yes | Yes | Alpha | Alpha | Alpha |
| Phylogenetic analysis | Yes | Yes | Yes | Yes | No | Tree |
| Feature abundance plots | Yes | Yes | Yes | Yes | Yes | Yes |
| Ordination plots | Yes | Yes | Yes | Yes | Yes | Yes |
| Network plots | Yes | No | No | Yes | No | Yes |
| Geographic distribution plots | No | No | No | No | No | Yes |
| Statistics plots | Yes | No | NR | Yes | NR | NR |
| Interactive visualization | 31 | 2;P | 3 | 8 | 9 | 17 |
| Raw data storage | No | No | Yes | No | No | Yes |
| Result storage | Yes | No | Yes | No | No | Yes |
| Online web Interface | Yes | No | Yes | No | Yes | Yes |
| R packaging | No | NR | NR | Yes | Yes | NR |
| Docker | Yes | No | No | No | Yes | No |
| Conda | Yes | Yes | Yes | No | Yes | No |

D: Export from DESeq2, M: Export from MetagenomeSeq, NR: Non relevant feature, P: Import/Export to Phyloseq, Number of unique interactive visualization are reported for each application in section 'Interactive visualization'

### User case

To illustrate how SHAMAN works, we performed the analysis of two sequencing experiments: a mock sequencing and a published dataset, afribiota dataset(48). In both analyses, we submitted the raw FASTQ files and provided a target file containing sample information (needed for statistical analysis).

### Zymo Mock dataset

The mock sequencing (EBI ENA code PRJEB33737) of the ZymoBIOMICS™ Microbial Community DNA was performed with an Illumina MiSeq. The Zymo mock community is composed with 8 phylogenetically distant bacterial strains, 3 of which are gram-negative and 5 of which are grampositive. DNA of two yeast strains that are normally present in this community were not amplified. Genomic DNA from each bacterial strain was mixed in equimolar proportions (https://www.zymoresearch.com/zymobiomics-community-standard). We compared the impact of both the number of amplification cycles (25 and 30 cycles) and the amount of DNA loaded in the flow cell (0.5ng and 1ng), on the microbial abundance. Each sample was sequenced 3 times (experimental plan provided in supplementary materials). Sequencing report provided by the sequencing facilities indicated the presence of contaminants. To handle this issue, we filtered out the genera occurring in less than 12 samples and outliers with a reduced log abundance as compared to the other genera (Fig. S3). This process selected the 8 bacterial stains of Zymo mock (Fig. 1). We then defined a statistical model that included DNA amount and the number of amplification cycle as main effects and an interaction between these variables. The statistical comparison showed a significant impact of the number of amplification cycle compared to DNA amount. We found no differential features between 0.5 ng and 1 ng DNA for each possible number of cycle (25 and 30 cycles), while the comparison of number of amplification cycle for each given amount of DNA showed significant impact on the abundance of mock bacteria (Table. S1, S2). These results are in agreement with previous studies that presented the PCR-induced bias on equivalent mix(49, 50).

**Fig. 1.**
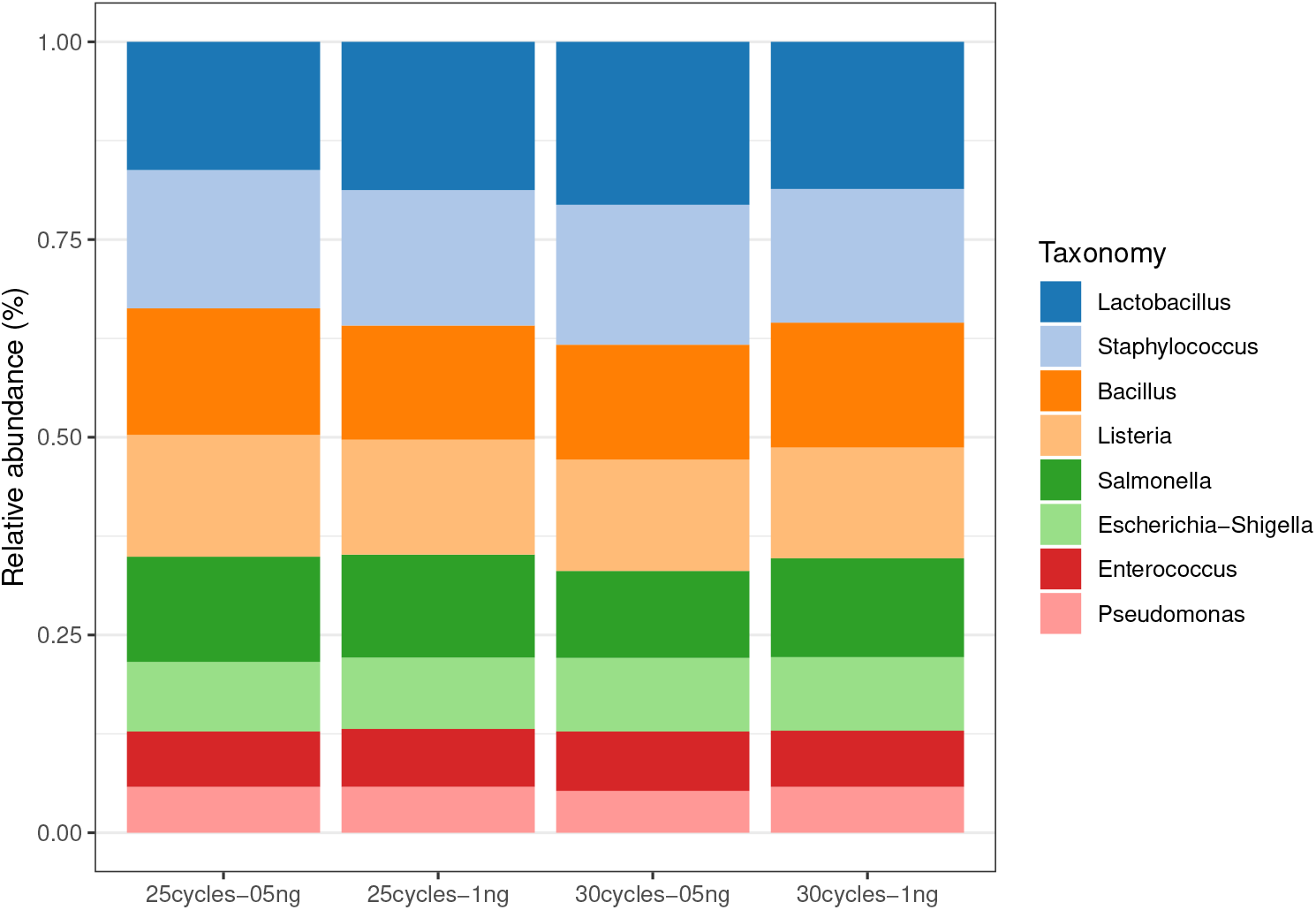
Barplot of taxa abundance of ZYMO MOCK samples. We summed the abundance of the OTU annotated at genera level with SILVA database and plotted the average abundance per condition.

### Afribiota dataset

The second dataset included samples of microbial communities in stunted children aged 2-5y living in sub-Saharan Africa (48). Three groups (nutritional status) of individuals were considered: NN=non stunted, MCM=moderately stunted, MCS=severely stunted. Samples originated from the small intestine fluids (gastric and duodenal) and feces. The authors performed the bioinformatic treatment with QIIME framework and the statistical analysis with several R packages including Phyloseq for the normalization and DESeq2 for the differential analysis. 541 samples were available on EBI ENA (code PRJEB27868).

Using SHAMAN, raw reads were filtered against Human HG38 and PhiX174 genomes. A total of 2386 OTUs were calculated and 76% were annotated with SILVA database at genus level. The sparsity rate of the contingency table was high with 0.84. In consequence, we used the weighted nonnull normalization which is particularly efficient when the matrix highly sparse (Fig. S2).

Two analyses were performed, a global analysis that included duodenal, gastric as well as feces samples and a more specific analysis including fecal samples only. Statistical models included the following variables, age, gender, country of origin and nutritional status. Overall our results obtained when using SHAMAN were highly consistent with those of Vonaesch et al. (48). We detected a significant change in the community composition between gastric and duodenal samples compared to feces samples at Genus level (Fig. 2a) (PERMANOVA, P=0.001). The most abundant genera were reported in Fig. S4. *α*-Diversity was not affected by stunting (Fig.2b). We looked for a distinct signature of stunting in the feces. We report in the volcano plot (Fig.2c) genera with differential abundance between stunt samples compared to non-stunt (complete list available in Table S3). Twelve microbial taxa, corresponding to members of the oropharyngeal core microbiota, were overrepresented in feces samples of stunted children as compared with non-stunted children; more particularly Porphyromonas, Neisseira and Lactobacillus (Fig.2d). These findings were in agreement with the conclusions of the Afribiota consortium while being obtained within a few minutes of interaction with the SHAMAN interface.

**Fig. 2.**
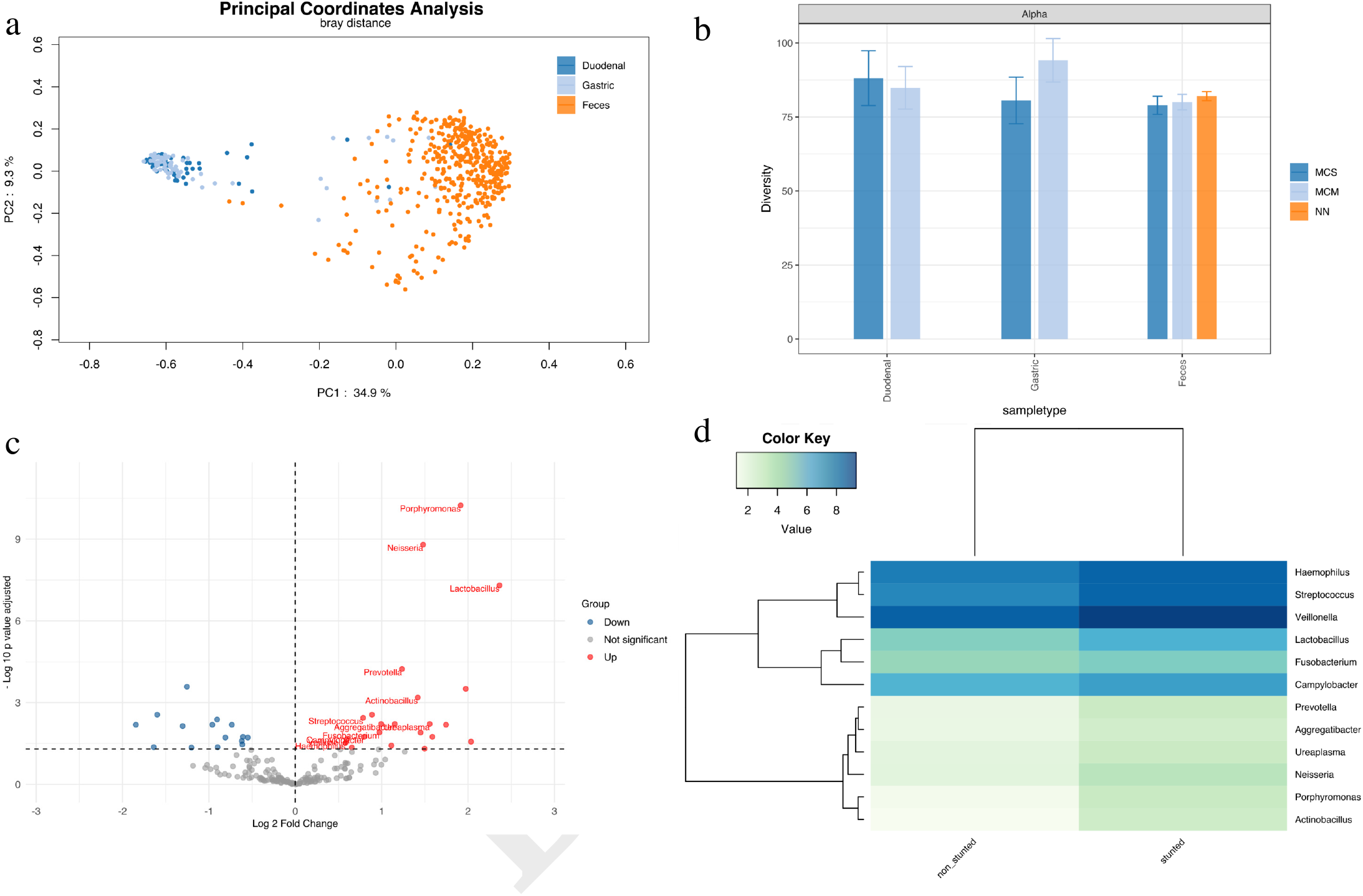
Afribiota study of small intestine fluids and feces from stunt children compared to non stunt. (a) PCoA plot the Bray-Curtis dissimilarity index of the samples. Duodenal samples are colored in blue, light blue for Gastric and orange for Feces. PERMANOVA test based on the sample type yielded a P value of 0.001. (b) Alpha diversity analysis of non-stunt (NN), moderately stunted (MCM) and severely stunted (MCS). Overlapping confidence interval indicates that the diversity are not different between NN, MCM and MCS in duodenal, gastric and feces samples. (c) Volcano plot of differentially abundant genera in the feces of stunt children compared to non-stunt. We plot the log2 fold change against the −log 10 adjusted p-value. Microbial taxa in red correspond to an increase of abundance and in blue to a decrease abundance. Labeled dots correspond to taxa from orpharyngeal core microbiota. (d) Log 2 abundance of differential abundant taxa from orpharyngeal core microbiota in stunt and non-stunt children feces.

## Conclusion

SHAMAN enables user to run most of the classical metagenomics methods and makes use of statistical analyses to provide support to each visualization. The possibility to deploy SHAMAN locally constitutes an important feature when the data cannot be submitted on servers for privacy issues or insufficient internet access. SHAMAN also simplifies the access to open computational facilities, making a careful use of the dedicated server, galaxy.pasteur.fr.

During its development, we felt a strong interest of the metagenomics community. We recorded 82 active users per month in 2019 (535 unique visitors in total) and 800 downloads of the docker application. We expect that SHAMAN will help researcher performing a quantitative analysis of metagenomics data.

## Data availability

Sequence reads of Zymo Mock have been deposited in the European Nucleotide Archive, https://www.ebi.ac.uk/ena/ accesion no. PRJEB33737.

## Acknowledgments

We thank Pascal Campagne for his comments, Hugo Varet for helpful discussions about DESeq2, Fabien Mareuil for the help to deploy SHAMAN computation on Galaxy and Youssef Ghorbal for the maintenance of the databank, as well as the IT System Department of Institut Pasteur, who manages installation and update of tools on TARS cluster.

