## Supplementary files for "SHAMAN: a user-friendly website for metataxonomic analysis from raw reads to statistical analysis"

### Appendix 1: Mathematical definition of the contingency table filtering

Let us denote by $\mathscr{F}$ the entire set of OTU. We propose to only consider a subset $\mathscr{R}$ of OTUs, for the analysis: $\mathscr{R} = \mathscr{L}_1 \cap \mathscr{L}_2$, where $\mathscr{L}_1$ is defined by

$$\mathscr{L}_1 = \left\{ f \in \mathscr{F} \middle| \sum_j 1_{\{c_{fj}>0\}} \geq l_1 \right\} \quad \text{with} \quad l_1 = \lfloor 0.8 \times k_{max} \rfloor,$$

where $c_{fj}$ is the abundance of the feature $f$ in the sample $j$ while $k_{max}$ is the maximum number of samples in which the feature is found. The subset $\mathscr{L}_2$ corresponds to the features with a not too small abundance and is defined by

$$\mathscr{L}_2 = \left\{ f \in \mathscr{F} \middle| \sum_j c_{fj} \geq l_2 \right\},$$

where $l_2$ is the intercept of the linear regression between the variables $y_k = \sum_i 1_{\{\sum_j c_{ij} > x_k\}}$ and $x_k \in \left[ \min_{f \in \mathscr{F}} \left( \sum_j c_{ij} \right) ; \lambda \right]$.

$\lambda$ is a tuning parameter whose default value is $\left\lfloor \frac{\sum_{ij} c_{ij}}{n} \times 0.05 \right\rfloor$.

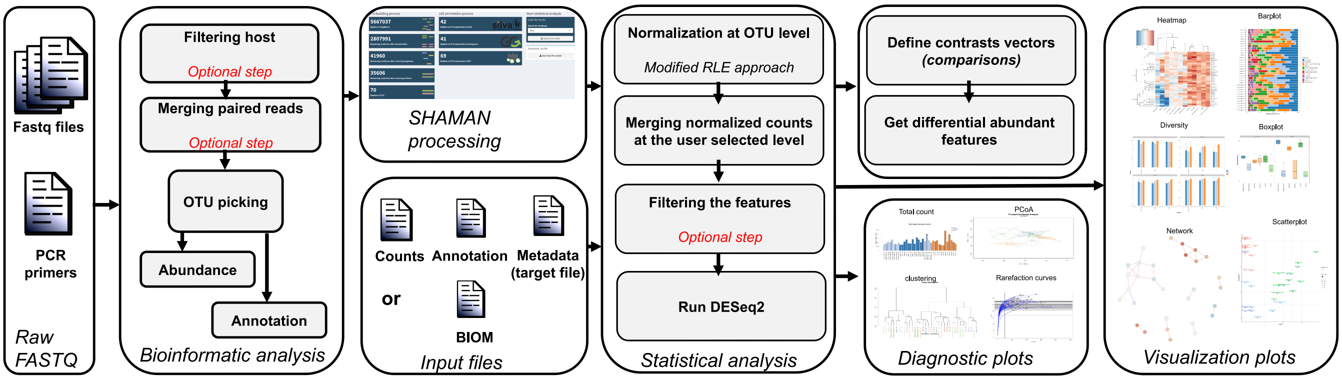


Figure 1: **Shaman workflow**. SHAMAN can start from raw reads or from processed data. In this last case, it needs at least three tables, the count matrix, the annotation table and the metadata which can either be provided into three different files or by using the BIOM format. The user can then select the variables of interest and add some batch effects. Once run, contrast vectors can easily be defined and interactive visualizations are available.

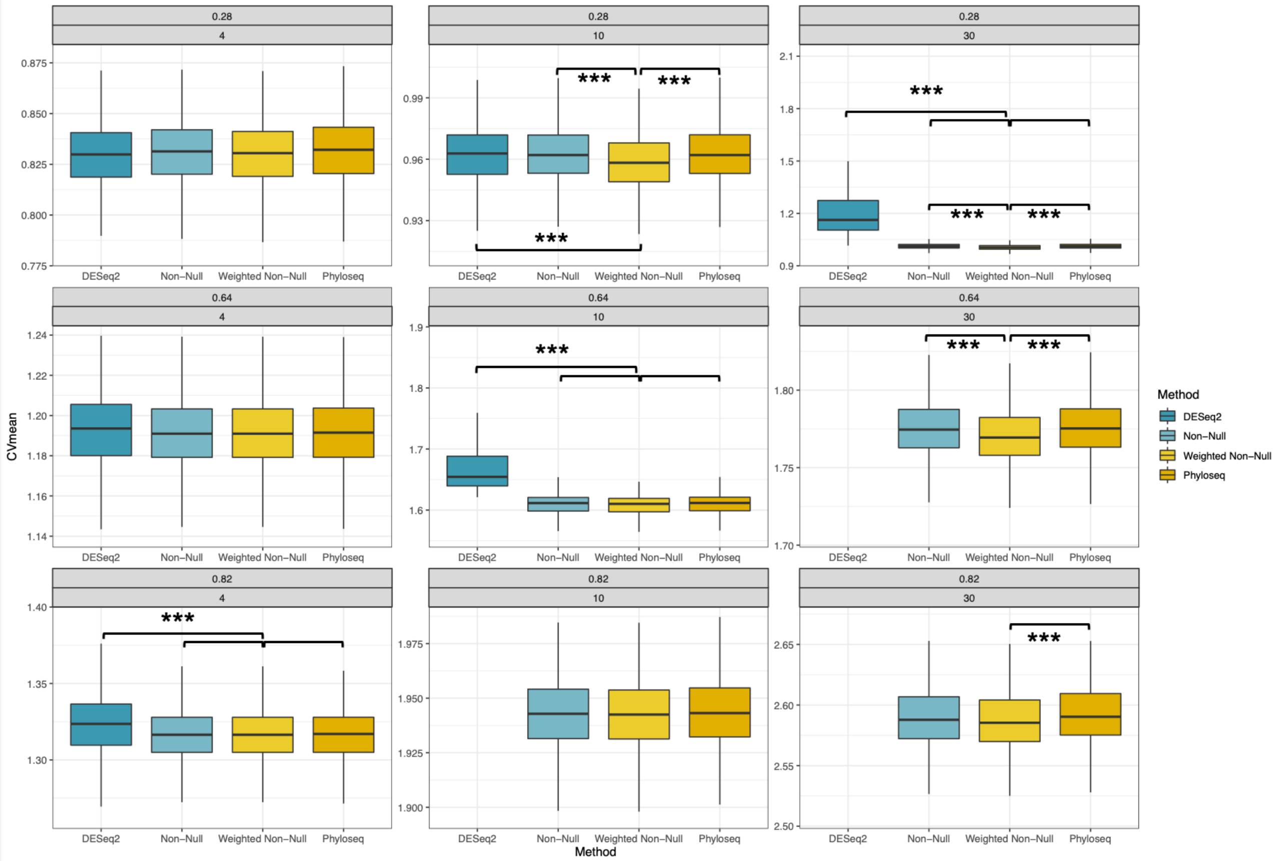


Figure 2: **Simulation study**. Boxplots of the average coefficient of variation for DESeq2, non-null, weighted non-null and Phyloseq normalization. The sparsity of the count matrices is chosen within $\{0.28, 0.64, 0.82\}$ and the number of observations vary within $\{4, 10, 30\}$. 500 normalizations were performed at each level of sparsity and for each number of observations. The results were analyzed by using a t-test, $^{***}p < 0.001$. DESeq2 normalization did not converged when the matrix sparsity and the number of observations was high (e.g., with a sparsity of 0.64 and 30 observations).

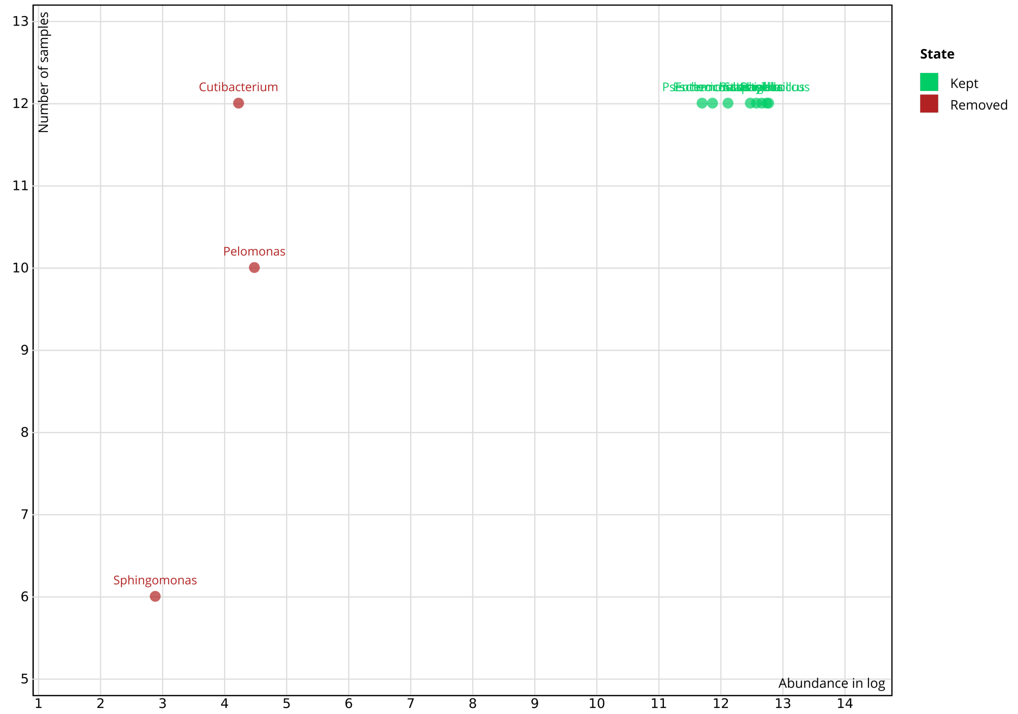


Figure 3: **Filtering of Zymo mock in SHAMAN**. Genera occurring in at least 12 samples and with log abundance $\geq 5$ were kept in the study (green dots). Cutibacterium, Pelomonas and Sphingomonas genera counts (red dots) were removed from the contingency table.

| ID | Base mean | Fold Change | Log2 fold change | P-value adjusted |
|---|---|---|---|---|
| Bacillus | 23314 | 1.613 | 0.69 | 0.0.0058 |
| Listeria | 22410 | 1.633 | 0.708 | 0.0.0058 |
| Salmonella | 19308 | 1.767 | 0.822 | 0.0.0058 |
| Pseudomonas | 8696 | 1.609 | 0.686 | 0.0045 |
| Staphylococcus | 26546 | 1.496 | 0.58 | 0.0045 |
| Escherichia-Shigella | 13921 | 1.432 | 0.518 | 0.0051 |
| Enterococus | 11024 | 1.417 | 0.504 | 0.0084 |

Table 1: **Summary of taxa differentially abundant when compared 25 and 30 amplification cycles for 0.5ng DNA load** Positive fold change indicates an increase of abundance of the taxa for the 25 amplification cycles.

| Id | Base mean | Fold change | Log2 fold change | P-value adjusted |
|---|---|---|---|---|
| Enterococcus | 11024 | 1.385 | 0.47 | 0.0303 |
| Listeria | 22410 | 1.427 | 0.514 | 0.0303 |
| Salmonella | 19308 | 1.473 | 0.559 | 0.0303 |
| Staphylococcus | 26546 | 1.381 | 0.465 | 0.0303 |
| Escherichia-Shigella | 13921 | 1.322 | 0.402 | 0.0393 |
| Pseudomonas | 8696 | 1.409 | 0.495 | 0.0414 |

Table 2: **Summary of taxa differentially abundant when compared 25 and 30 amplification cycles for 1ng DNA load** Positive fold change indicates an increase of abundance of the taxa for the 25 amplification cycles.

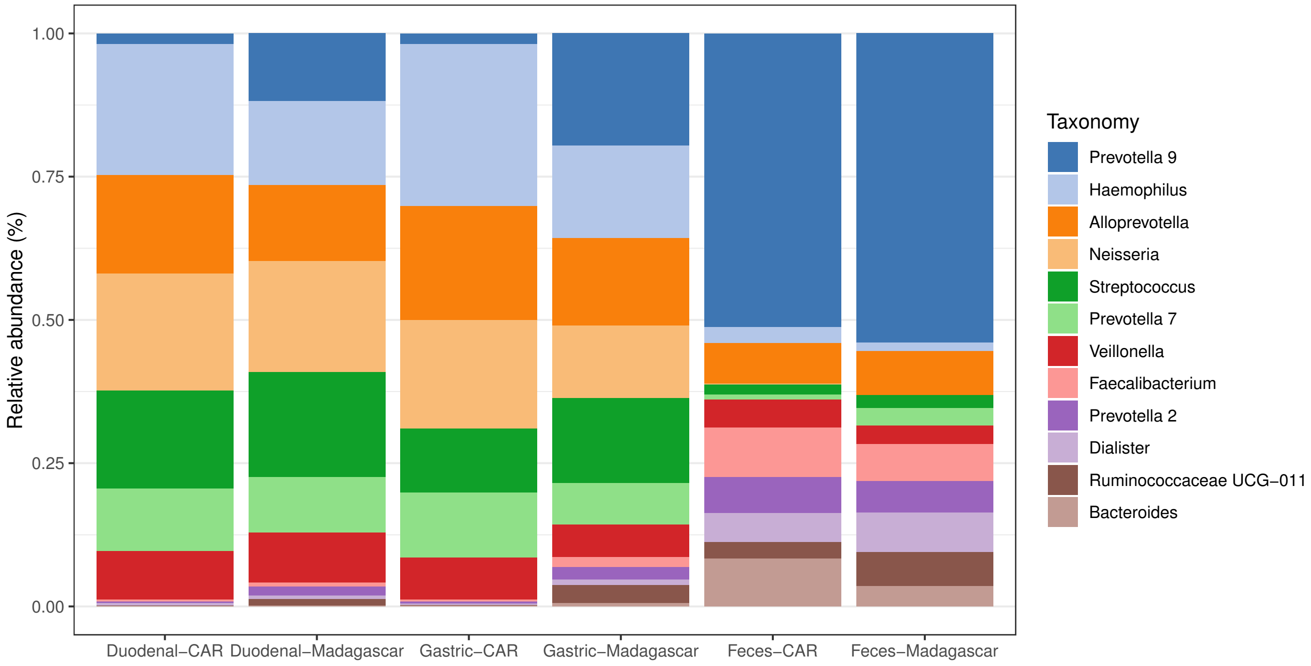


Figure 4: **Barplot of the 12 most abundant genera in duodenal, gastric and feces from children of Central African Republic (CAR) and Madagascar.** Haemophilus and Neisseria genera were also observed in cultured samples.

| Id | Base mean | Fold change | Log2 fold change | P-value adjusted |
|---|---|---|---|---|
| Porphyromonas | 4.73 | 3.772 | 1.915 | 5.755e-11 |
| Neisseria | 7.16 | 2.789 | 1.48 | 1.605e-09 |
| Lactobacillus | 51.69 | 5.152 | 2.365 | 4.962e-08 |
| Prevotella | 5.04 | 2.354 | 1.235 | 5.836e-05 |
| Ruminococcaceae UCG-009 | 3.46 | 0.419 | -1.255 | 0.0003 |
| Weissella | 11.33 | 3.927 | 1.974 | 0.0003 |
| Actinobacillus | 3.98 | 2.672 | 1.418 | 0.0006 |
| Rikenellaceae RC9 gut group | 86.21 | 0.33 | -1.598 | 0.0028 |
| Ruminococcaceae UCG-011 | 502.47 | 1.85 | 0.887 | 0.0028 |
| Streptococcus | 237.2 | 1.723 | 0.785 | 0.0036 |
| Christensenellaceae R-7 group | 49.22 | 0.534 | -0.906 | 0.0042 |
| Aggregatibacter | 4.31 | 2.222 | 1.152 | 0.0061 |
| Granulicatella | 0.81 | 1.991 | 0.993 | 0.0061 |
| Ureaplasma | 5.52 | 2.94 | 1.556 | 0.0061 |
| Ruminococcaceae UCG-002 | 144.01 | 0.601 | -0.735 | 0.0065 |
| Streptobacillus | 0.63 | 3.351 | 1.745 | 0.0065 |
| Terrisporobacter | 4.29 | 0.514 | -0.961 | 0.0065 |
| [Eubacterium] xylanophilum group | 2.05 | 0.405 | -1.305 | 0.0073 |
| Fusobacterium | 28.55 | 1.964 | 0.974 | 0.0124 |
| Abiotrophia | 0.45 | 2.733 | 1.45 | 0.0126 |
| Campylobacter | 89.02 | 1.739 | 0.798 | 0.0181 |
| [Eubacterium] coprostanoligenes group | 119.89 | 0.657 | -0.606 | 0.0181 |
| Capnocytophaga | 0.19 | 3.005 | 1.587 | 0.0181 |
| Ruminococcaceae UCG-005 | 211.21 | 0.683 | -0.55 | 0.0192 |
| Ruminococcaceae UCG-010 | 31.58 | 0.57 | -0.81 | 0.0192 |
| Veillonella | 475.74 | 1.515 | 0.599 | 0.0218 |
| Ruminococcaceae NK4A214 group | 36.02 | 0.651 | -0.62 | 0.0257 |
| Morganella | 0.42 | 4.099 | 2.035 | 0.0271 |
| Haemophilus | 260.89 | 1.494 | 0.58 | 0.0301 |
| Family XIII AD3011 group | 9.67 | 0.653 | -0.614 | 0.0344 |
| Lactococcus | 35.73 | 2.161 | 1.112 | 0.0373 |
| Methanobrevibacter | 5.99 | 0.321 | -1.638 | 0.0429 |
| Turicibacter | 10.03 | 0.534 | -0.9 | 0.0429 |
| Escherichia-Shigella | 25.03 | 1.574 | 0.654 | 0.0444 |
| Kingella | 0.47 | 2.822 | 1.497 | 0.0488 |

Table 3: **Summary of taxa differentially abundant when compared samples of stunted to non stunted children.** Positive and negative fold changes indicate respectively an increase of abundance of the taxa in stunted and non-stunted children.
